# Sex-biased long non-coding RNAs negatively correlated with sex-opposite protein coding gene co-expression networks in Diversity Outbred mouse liver

**DOI:** 10.1101/271668

**Authors:** Tisha Melia, David J. Waxman

**Author notes:** Correspondence: Dr. David J. Waxman, Dept. of Biology, Boston University, 5 Cummington Mall, Boston, MA 02215.

## Abstract

Sex differences in liver gene expression and disease susceptibility are regulated by pituitary growth hormone secretion patterns, which activate sex-dependent liver transcription factors and establish sex-specific chromatin states. Ablation of pituitary hormone by hypophysectomy (hypox) has identified two major classes of liver sex-biased genes, defined by their sex-dependent positive or negative responses to hypox, respectively; however, the mechanisms that determine the hypox responsiveness of each gene class are unknown. Here, we sought to discover candidate regulatory long noncoding RNAs (lncRNAs) that control hypox responsiveness. First, we used mouse liver RNA-seq data for 30 different biological conditions to discover gene structures and expression patterns for ~15,500 liver-expressed lncRNAs, including antisense and intragenic lncRNAs, as well as lncRNAs that overlap active enhancers, marked by enhancer RNAs. We identified >200 robust sex-specific liver lncRNAs, including 157 whose expression is regulated during postnatal liver development or is subject to circadian oscillations. Next, we utilized the high natural allelic variance of Diversity Outbred (DO) mice, a multi-parental outbred population, to discover tightly co-expressed clusters of sex-specific protein-coding genes (gene modules) in male liver, and separately, in female liver. Sex differences in the gene modules identified were extensive. Remarkably, many gene modules were strongly enriched for male-specific or female-specific genes belonging to a single hypox-response classes, indicating that the genetic heterogeneity of DO mice captures responsiveness to hypox. Hypox-responsiveness was shown to be facilitated by multiple, distinct gene regulatory mechanisms, indicating its complex nature. Further, we identified 16 sex-specific lncRNAs whose expression across DO mouse livers showed an unexpected significant negative correlation with protein-coding gene modules enriched for genes of the opposite-sex bias and inverse hypox response class, indicating strong negative regulatory potential for these lncRNAs. Thus, we used a genetically diverse outbred mouse population to discover tightly co-expressed sex-specific gene modules that reveal broad characteristics of gene regulation related to responsiveness to hypox, and generated testable hypotheses for regulatory roles of sex-biased liver lncRNAs that control the sex-bias in liver gene expression.

## Introduction

Sex differences in hepatic gene expression are extensive [1–3], and are associated with sex differences in liver function and disease susceptibility [4–8]. These sex differences are primarily regulated by growth hormone (GH) [9–11], working through its sex-differential pituitary secretory patterns: pulsatile in male and near-continuous in female [12–15]. The resultant sex-differential plasma GH patterns regulate the activity of STAT5 and downstream transcription factors in a sex differential manner [14–18]. These transcription factors are essential for the complex transcriptomic patterns specific to each sex [1, 19] and help establish sex-differences in chromatin accessibility and chromatin states closely linked to sex-specific gene expression [20, 21]. The sex-biased transcriptome includes long noncoding RNAs (lncRNAs) [22], which have the potential for diverse regulatory roles, including epigenetic and transcriptional regulation of both in *cis* and in *trans* [23–28].

Studies in rodent models have established that the prominent, sex-biased temporal patterns of GH secretion, are imprinted in males by neonatal exposure to androgens [29]. Sex-specific liver gene expression does not, however, become widespread until puberty [30, 31]. Surgical removal of the pituitary gland by hypophysectomy (hypox) in mice, and rats, identifies two major classes of sex-biased genes in each sex, based on their responses to the loss of GH and other pituitary hormones [32, 33]. Male class 1 and female class 1 liver sex-specific genes are activated by the GH secretion pattern of the sex where they are more highly expressed, whereas, male class 2 and female class 2 liver sex-specific genes are repressed by the GH secretion pattern of the sex where they are less highly expressed (Fig. S1). Consequently, following hypox, class 1 genes are down regulated and class 2 genes are de-repressed (up regulated) in mouse liver, which leads to a near global loss of sex-specific gene expression [32, 33]. The molecular basis for these distinct hypox response phenotypes is poorly understood.

Gene clustering has been used to identify groups of genes (gene modules/gene clusters) whose expression is strongly associated with different molecular phenotypes (gene signatures), including cancer subtypes [34–37] and various cellular processes [38–40]. An early study used gene modules discovered in mouse fat, brain, liver and muscle to study sex differences in the organization of gene expression networks. Gene modules discovered in liver and fat were found to be the least conserved between the sexes [41].

Here, we use mouse liver RNA-Seq data for 30 distinct biological conditions to discover and characterize more than 15,000 liver-expressed lncRNAs, including many sex-dependent lncRNAs subject to postnatal regulation during liver development [31, 42] or circadian oscillations [43, 44]. We exploited the high genetic variability of DO mice, a multi-parental outbred population derived from eight genetically divergent inbred mouse strains [45, 46], to discover biologically meaningful clusters of sex-biased genes, and we investigated their enrichment for each of the above hypox-response classes. Our findings reveal a striking, and unexpected, significant negative correlation across a large panel of DO mouse livers between sex-specific lncRNAs and gene modules enriched for protein-coding genes of the opposite sex and inverse hypox-response class, generating testable hypotheses for regulatory roles of sex-biased liver lncRNAs.

## Materials and Methods

### LncRNA discovery

We collected 186 RNA-Seq samples of male and female mouse liver, representing 30 different biological conditions (Table S1A) [31, 32, 43, 47–50]. Tophat2 [51] was used to map a total of 10.7 billion reads to the C57Bl/6J mouse mm9 reference genome. LncRNA gene structures and isoforms were discovered using the method we recently described [22], with the following changes: (1) the expression filter was adjusted to remove lowly expressed genes at the level of 10% false positive rate (FPR), instead of 15%; (2) lncRNA gene structures whose exons overlapped one or more protein-coding gene exons were removed only if they were on the same strand as the protein-coding gene. The latter modification allowed us to retain gene structures for antisense lncRNAs and intragenic lncRNAs. LncRNAs that did not overlap with any protein-coding gene were designated intergenic. LncRNAs whose exons overlap protein-coding gene introns were designated antisense if they were transcribed on the opposite strand as the protein-coding gene; otherwise they were designated intragenic.

### Gene expression quantification for lncRNAs

We devised four approaches for counting sequence reads to quantify lncRNA gene expression: (1) counting reads that overlap any exon; (2) counting reads that overlap any region from the transcription start site (TSS) to the transcript end site (TES) (gene body); (3) counting reads that overlap genomic regions that were assigned as exons in all isoforms of a gene (exon-only counting); and (4) counting reads that overlap genomic regions that were designated introns in all isoforms of a gene (intron-only counting). These counting methods closely follow a recent publication for protein-coding genes [32]. Reads that mapped to regions that can be attributed to more than one gene were not counted, consequently, intragenic lncRNAs that are wholly within another gene will have no read counts when using counting methods 2 and 4. All of the analyses presented in this study are based on the first counting method.

### Sex-specific and GH-regulated lncRNAs

Eight mouse liver RNA-seq datasets (Table S1B) were used to assess the sex-specificity of lncRNA gene expression, as follows. Datasets # 1-3: Three independent poly(A)-selected CD-1 mouse total liver RNA datasets, which together encompassed 7 pools from each sex [32, 48]. Datasets # 4-6: For each sex, three pools of poly(A)-selected nuclear RNA, three pools of ribosomal RNA-depleted total RNA, and three pools of ribosomal RNA-depleted nuclear RNA, all from CD-1 mouse liver [48]. Dataset # 7: 12 male and 12 female livers from 15-20 week old C57Bl/6J mouse liver [31]. Dataset # 8: 12 male and 12 female C57Bl/6J mouse livers collected on embryonic day 17.5 [31]. Differential analysis using edgeR [52] was performed to compare RNA expression levels in male *vs* female livers for each of the eight datasets. Any lncRNA with a male/female expression ratio (|fold-change| > 4 at FDR < 0.05) in any of the eight datasets was designated as a sex-specific lncRNA. A listing of all 684 sex-specific lncRNAs that met these criteria for one or more of the 8 datasets is provided in Table S2A, along with their expression values.

### Pituitary regulation

Pituitary-regulated lncRNAs are those that showed a significant response to hypox in either male or female liver at |fold-change| > 2 and FDR < 0.05 for comparisons between hypox and intact mice, as determined by edgeR. Sex-specific liver lncRNAs were categorized into four classes based on their response to hypox: class 1 and class 2 male-specific lncRNAs, and class 1 and class 2 female-specific lncRNAs. RNA-seq data from two independent hypox mouse liver datasets were used to establish each lncRNA’s response to hypox: PolyA-selected liver RNA and ribosomal-depleted liver RNA [32]. Initial analysis of the ribosomal RNA-depleted hypox liver RNA dataset was carried out by Christine Nykyforchyn of this lab. All but four lncRNAs were assigned to the same hypox-response class by both hypox datasets; those four inconsistencies were resolved by choosing the classification indicated by the dataset with the higher male/female |fold-change|. The hypox-response class of each sex-specific lncRNA is shown in Table S2A.

### Regulated lncRNAs during liver development

edgeR was used in ten separate differential expression analyses of C57BL/6J male mouse liver RNA-Seq datasets comparing lncRNA expression profiles between day 0 (immediately after birth and before the start of suckling) with liver RNA-Seq datasets for each of the following postnatal ages: day 1 (exactly 24 h after birth) and days 3, 5, 10, 15, 20, 25, 30, 45 and 60 [52]. LncRNAs with gene expression |fold-change| > 2 at FDR < 0.05 at any of the above ages were designated developmentally regulated lncRNAs. See Table S2A for a listing of developmental data for liver lncRNAs.

### Regulated lncRNAs in a 24-hour period

Five differential expression analyses were performed to compare liver lncRNA RNA-seq expression profiles in C57BL/6J male liver at Zeitgeber times (ZT) ZT6, ZT10, ZT14, ZT18, ZT22 to gene expression at ZT2 using edgeR. LncRNAs with gene expression |fold-change| > 2 at FDR < 0.05 at any ZT time point were designated circadian regulated lncRNAs. Circadian expression data for liver lncRNAs is shown in Table S2B.

### Gene expression quantification in Diversity Outbred (DO) mouse liver

RNA-Seq data obtained for comprised of 112 male and 121 female DO mouse livers samples, all from mice fed standard chow diet, were downloaded from GEO, accession numbers GSE45684 [53, 54] and GSE72759 [55]. Read mapping and gene expression quantification for each sample was performed as follows. Briefly, reads were mapped to a specific diploid genome constructed for each individual mouse. Gene expression was then quantified by counting sequenced reads that overlap any exon by at least one bp using featureCounts [56], using the best mapped location for each read. The restriction of only using unique reads for diploid genomes would limit the read counts to only include those reads that mapped to genome locations where the paternal and maternal allele differ, and thus could substantially underestimate the level of expression of any given gene. The gene expression level obtained for diploid genomes is based on the total number of reads that overlap the counted regions in either the paternal or maternal allele. Reads counts were then transformed to fragments per kilobase of exon per million reads mapped (FPKM) for downstream analysis, where gene lengths are the average length of the gene in the two alleles. A batch effect that correlated with the GEO accession of the samples was removed from the gene expression level (transformed to log2(FPKM + 1)) using the ComBat function in the sva R package [57]. A listing of expression level across the full set of male and female DO mouse livers is provided in Table S7.

### Gene expression quantification for antisense lncRNAs based on unstranded RNA-Seq datasets

The FDR for antisense lncRNAs was set to 1 for all differential analyses performed using unstranded RNA-seq datasets, as the unstranded nature of such data prevents us from reliably attributing sequence reads to antisense lncRNAs. Unstranded RNA-seq datasets include: the first 2 datasets used to determine lncRNAs sex-specificity (Table S1B; *first two rows*), the polyA-selected hypox dataset used to identify pituitary hormone-regulated lncRNAs (Table S1A; *rows: male hypox and female hypox*), liver development RNA-seq datasets in male liver (Table S1A; *any row with samples from* [49]), RNA-seq datasets for DO mouse liver samples, and RNA-seq datasets that assess liver gene expression changes due to circadian rhythm (Table S1A; *rows: Male circadian ZT2-ZT22*).

### Sex-specific and GH-regulated protein-coding genes

We used a set of 1,033 sex-specific genes protein coding, which consists of genes that showed significant sex-specific gene expression in either CD-1 mouse liver or in any of the eight DO mouse founder strains (A/J, C57BL/6J, 129S1/SvlmJ, NOD/ShiLtJ, NZO/HILtJ, CAST/EiJ, PWK/PhJ and WSB/EiJ) [45, 58]. A list of 531 pituitary hormone-regulated protein-coding genes, defined as genes responsive to hypox, was downloaded from Table S3 in [32].

### Clustering sex-specific protein-coding genes

Sex-specific protein-coding genes were clustered based on their gene-expression correlation across 112 male DO mouse livers, and separately across 121 female DO mouse livers, using the weighted correlation network analysis (WGCNA) [59]. WGCNA identifies co-expressed groups of genes (i.e., gene clusters, also referred to as gene modules or gene networks) based on pairwise gene expression correlations across all samples. The correlation matrix is transformed into an adjacency matrix, which reflects the distance between genes, based on weighted gene expression correlations. Weights are introduced by raising the actual gene expression correlation to a power of β, which leads to more robust gene similarity measures. WGCNA parameters were set as follows: correlation function = the bicor function [60] with the maxPOutliers set to 0.1, type of network = signed hybrid, minimum module size = 5 genes, soft thresholding power (ß) = 3, deepSplit = 3 and cutTreeHeight = 0.25 to merge similar modules. Other parameters were used at their default settings. Genes that were not expressed (FPKM = 0) in any set of livers (male liver samples, or separately, female liver samples) were excluded from the clustering analysis, as were genes whose variance is 0 (MAD = 0). This resulted in 1,018 and 1,014 of the 1,033 sex-specific protein-coding genes (Table S3-1) being clustered in male and female DO mouse liver samples, respectively; the overlap between the two gene sets was high (1,013 of 1,019 genes).

### Relationship between protein-coding gene modules

Similarities between protein-coding gene modules in each sex were measured by pairwise Pearson correlation between their first principal components, as calculated using the moduleEigenGenes function in WGCNA. The correlation matrix was then clustered using average linkage hierarchical clustering with default parameters.

### Enrichment of various gene sets for protein-coding gene clusters (gene modules)

To determine if any of the identified protein-coding gene clusters is enriched for any gene set with biological meaningful properties, we used the Hypergeometric test to assess the significance of overlap between the gene sets being compared. An overlap at p < 0.05 indicates significant enrichment for the overlapping gene set. Enrichment for each of the following gene sets was tested for each gene cluster: male-specific genes, female-specific genes, strongly sex-specific genes (i.e., male/female |fold-change| > 4), male class 1 genes, male class 2 genes, female class 1 genes, and female class 2 genes (see Table S3-1 for the first three categories, and see Table S3 in [32] for members of the hypox-response gene classes). To calculate the enrichment of male-specific genes in cluster *i*, we let *mc* = number of male-specific genes in cluster *i*, *gc* = number of genes in cluster *i*, *g* = number of genes included in the cluster analysis (1,018 sex-specific genes in male liver samples, and 1,014 sex-specific genes in female liver samples, as described above) and *m* = the number of male-specific genes in *g*. Next, the probability of observing at least *x* male-specific genes in cluster *i* (P(#male-specific genes >= *x*)) was defined by the probability of drawing *x* or more male-specific genes, given that we are drawing *gc* number of genes from *g* genes, which has *m* male-specific genes. This was implemented using the following R command: 1 - phyper(*x*-1, *m*, *g-m*, *gc*, lower.tail = TRUE, log.p = FALSE). The same formula was used to calculate enrichment for other gene sets, except we replaced the male-specific gene set with the other gene sets of interest.

### Properties of protein-coding gene clusters

Nine properties were assessed for each gene cluster (see lower panels in Fig. 3 and Fig. 4): (1) the number of genes in each cluster; (2) the number of distinct TADs where the genes in each cluster are found; (3) the percentage of genes in each cluster that show strong sex-bias (male/female |fold-change| > 4 at FDR < 0.05; Table S3-1); the percentage of male-specific (4) or female-specific (5) genes in each cluster (Table S3-1); and the number of male-specific class 1 (6), male-specific class 2 (7), female-specific class 1 (8) and female-specific class 2 (9) genes in each cluster, expressed as a percentage of all hypox-responsive genes (Table S3 in [32]) in the cluster. The percentages in rows 6-9 of Fig. 3 and Fig. 4 add up to 100%. For properties # 3-9, percentage values are only shown for those gene clusters that show a significant overlap with the relevant gene set, i.e. p-value < 0.05, as determined by the Hypergeometric test, described above.

### Correlating lncRNA with protein-coding genes clusters

To identify lncRNAs that show a significant positive correction or a significant negative correlation with protein-coding gene clusters, we calculated the Pearson correlation between: 1) the expression pattern of each sex-specific lncRNA across the full set of 112 male DO mouse livers, and separately, across the full set of 121 female DO mouse livers; and 2) the first principal component of each protein-coding gene module across the same DO mouse liver sets. The first principal component of each protein-coding gene module was calculated using the moduleEigenGenes function in WGCNA with default parameters. We determined all lncRNA-gene module correlation pairs in male DO liver samples, and separately in female DO liver samples. P-values for the significance of the correlations were obtained using the cor.test function in R and adjusted by the FDR method. Due to the unstrandedness of the DO mouse liver RNA-seq datasets, this analysis was limited to the set of 168 multi-exonic, intergenic sex-specific lncRNAs (Table S2A). Top correlations were those that met a p-value of < 0.01 at FDR < 0.001.

### eRNA identification

27 GRO-seq C57BL/6J male liver samples were downloaded using GEO accession # GSE36916 and # GSE59486 [43, 44]. Poly-A tails were trimmed from sequence reads using HOMER [61] using the following parameters: motif: AAAAAA or TTTTTT, mismatch = 2, minMatchLength = 5 and-min = 20. Sequence reads were mapped to the C57BL/6J mm9 reference genome using Bowtie2 [62], and mapped reads were used to discover peaks using HOMER with the follower parameter settings: L = 3 and bodyFold = 3. Every peak in the positive strand was paired with the nearest peak in the reverse strand. Any peak pairs that were > 500 bases apart and converging into the same direction were removed. eRNA loci were defined as the 1 kb region centered at the overlapping region in each peak pair. The 15,853 eRNA loci that overlap any published H3K27ac or H3K4me1 peaks for young adult mouse liver [21] were retained (Table S6).

## Results

### Discovery of 15,558 liver-expressed lncRNAs

We recently described 4,961 mouse liver-expressed intergenic lncRNAs [22] and characterized their gene expression, conservation level and transcriptional regulation. We, and others, have since produced many strand-specific RNA-seq datasets, where the direction of transcription is preserved. These datasets enabled us to discover other classes of lncRNA that were not included in our initial study, namely antisense and intragenic lncRNAs, many of which may contribute to the transcriptional regulation of nearby genes [26, 27, 63]. We collected 186 RNA-seq datasets encompassing mouse livers representing 30 different biological states, spanning from embryonic day 15.5 to 15-20 week old mice (Fig. 1A). Other biological conditions represented include livers from each of the following: mice whose GH secretion patterns were altered, mice treated with foreign chemical agonists of liver nuclear receptors, and mice euthanized every 4 h to identify gene expression changes due to circadian rhythm [31, 32, 43, 47–50]. A total of 10.7 billion reads were mapped to the reference genome. LncRNAs were discovered using the methods described in [22], except that we only removed gene structures whose exons overlap protein-coding gene exons transcribed in the same direction (Fig. S2). 15,558 lncRNAs were discovered (Table S2A), of which 13,343 did not overlap any protein-coding gene, and were therefore designated intergenic (Fig. 1B). 1,966 lncRNAs were designated antisense lncRNAs based on their overlap with the antisense strand of protein-coding genes (Fig. 1C). 249 lncRNAs were intragenic, i.e., they were transcribed in the same direction as an overlapping protein-coding gene, but without any exonic region overlap (Fig. 1D). 16% (2,098) of the intergenic lncRNAs have multiple exons (multi-exonic lncRNAs), while 47% (916) of the antisense and 55% (138) of the intragenic lncRNAs are multi-exonic. A subset of lncRNAs (18%; 2,847) may be involved in the activation of other genes due to their overlap with enhancer RNAs (eRNAs) (Table S2A), which were previously shown to correlate positively with enhancer activity [64]. Overall, 44% of the 15,558 liver-expressed lncRNAs are not found in any prior annotation or database for noncoding RNAs [22, 65, 66].

**Figure 1.**
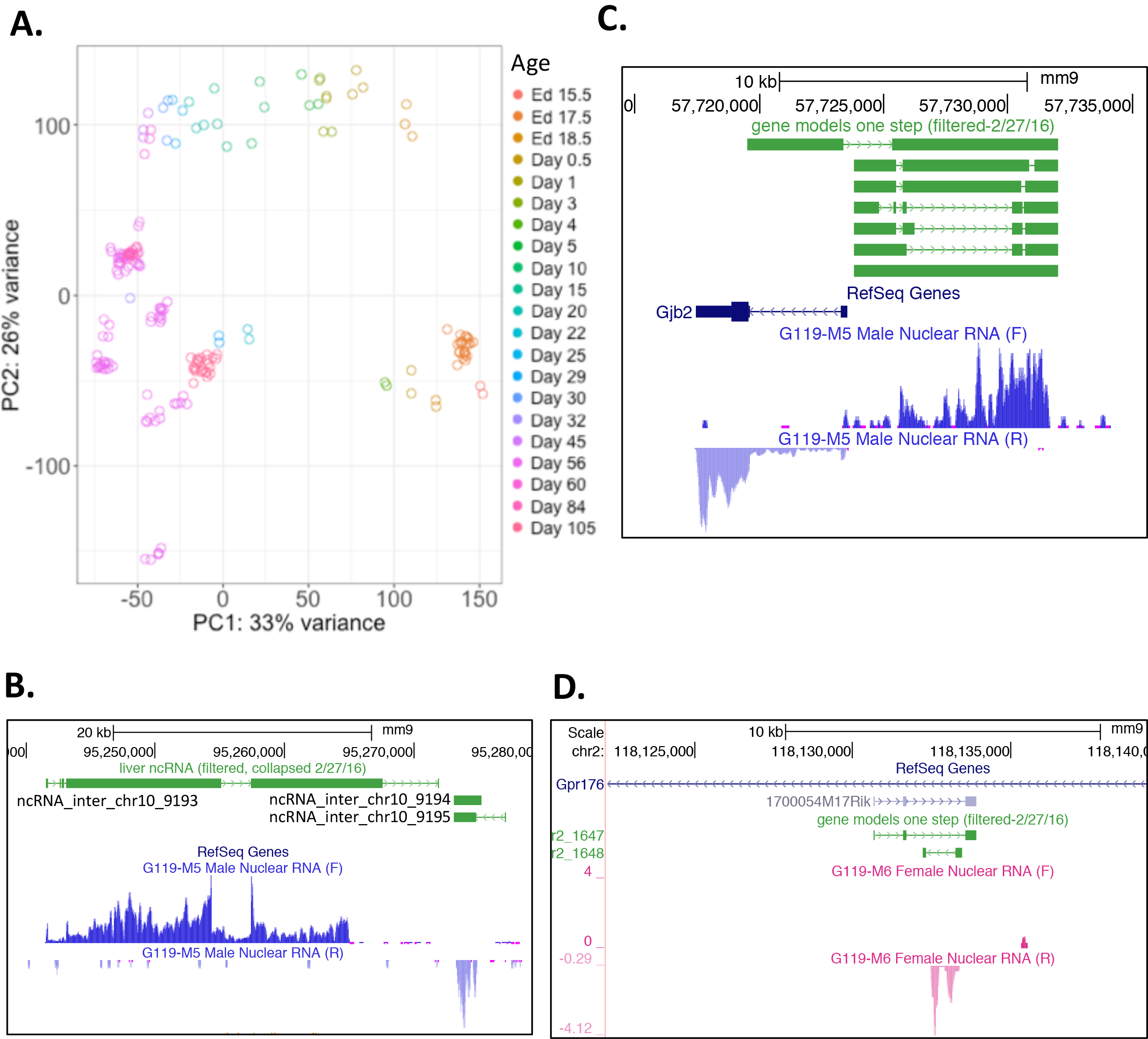
Liver-expressed lncRNAs. <u>**(A)**</u> PCA of the 186 RNA-Seq samples used to discover liver-expressed lncRNAs. The first principal component corresponds to the age of the mice from young to old (right to left). <u>**(B)**</u> Examples of novel intergenic lncRNAs discovered in this study: ncRNA_inter_chr10_9193, ncRNA_inter_chr10_9194 and ncRNA_inter_chr10_9195. Gene structures of lncRNAs are shown in green. Liver gene expression in select samples is shown as wig files. F, forward strand, R, reverse strand. <u>**(C)**</u> An antisense lncRNA, ncRNA_as_chr14_12073, that is transcribed on the forward strand and overlaps *Gjb2*, which is transcribed on the reverse strand. <u>**(D)**</u> An intragenic lncRNA, ncRNA_intra_chr2_1648, that is transcribed on the reverse strand, overlaps the gene *Gpr176* as well as another annotated lncRNA (1700054M17Rik).

### Regulation of sex-specific lncRNAs in male liver

The liver transcriptome undergoes significant changes during postnatal liver development [42, 67]. Notably, sex-biased gene expression emerges at ~4 weeks of age, with many more sex-specific genes showing developmentally regulated expression in male liver as compared to female liver [30]. We identified 684 sex-specific lncRNAs that showed significant sex-biased gene expression (male/female |fold-change| > 4 at FDR < 0.05; Table S2A) in any of 8 independent datasets comparing male liver to female liver samples (Fig. S3, *left panel*). 168 of these lncRNAs were multi-exonic and intergenic. We assessed their regulation during postnatal male liver development, by comparing their gene expression patterns on day 0 (immediately after birth and before the start of suckling) to that on day 1 (i.e., 24 h after birth), and to that on days 3, 5, 10, 15, 20, 25, 30, 45 and 60 of age (Fig. S3, *right panel*). 144 of the sex-specific lncRNAs were regulated at least 2-fold at FDR < 0.05 on at least one of the above developmental time points (Fig. 2A; Table S2A and S2C). The majority of male-biased lncRNAs were up regulated by 25 days of age in male liver, consistent with the patterns found for male-biased protein-coding genes (Fig. 2, *gene set I*) [30]. A subset of the male-specific genes in set I, however, showed a significant increase in expression as early as the first week of life. Correspondingly, a subset of female-specific genes was down regulated in male liver by 20 days of age (Fig 2, *set II*). A smaller subset of male-specific genes was strongly repressed after birth, and showed either continued repression (Fig. 2, *set III*) or up regulation around 30 days of age (Fig. 2, *set IV*), while maintaining their male-biased expression in adulthood. Most female-specific genes were up regulated earlier in male liver development and were either maintained or tapered off their up regulation at adulthood (60 days of age; Fig. 2, *set V*). The latter gene set still showed female-biased expression in adulthood, likely due to a stronger up regulation in female liver, as compared to male liver. This pattern was previously seen for a limited number of female-specific protein-coding genes [30].

**Figure 2.**
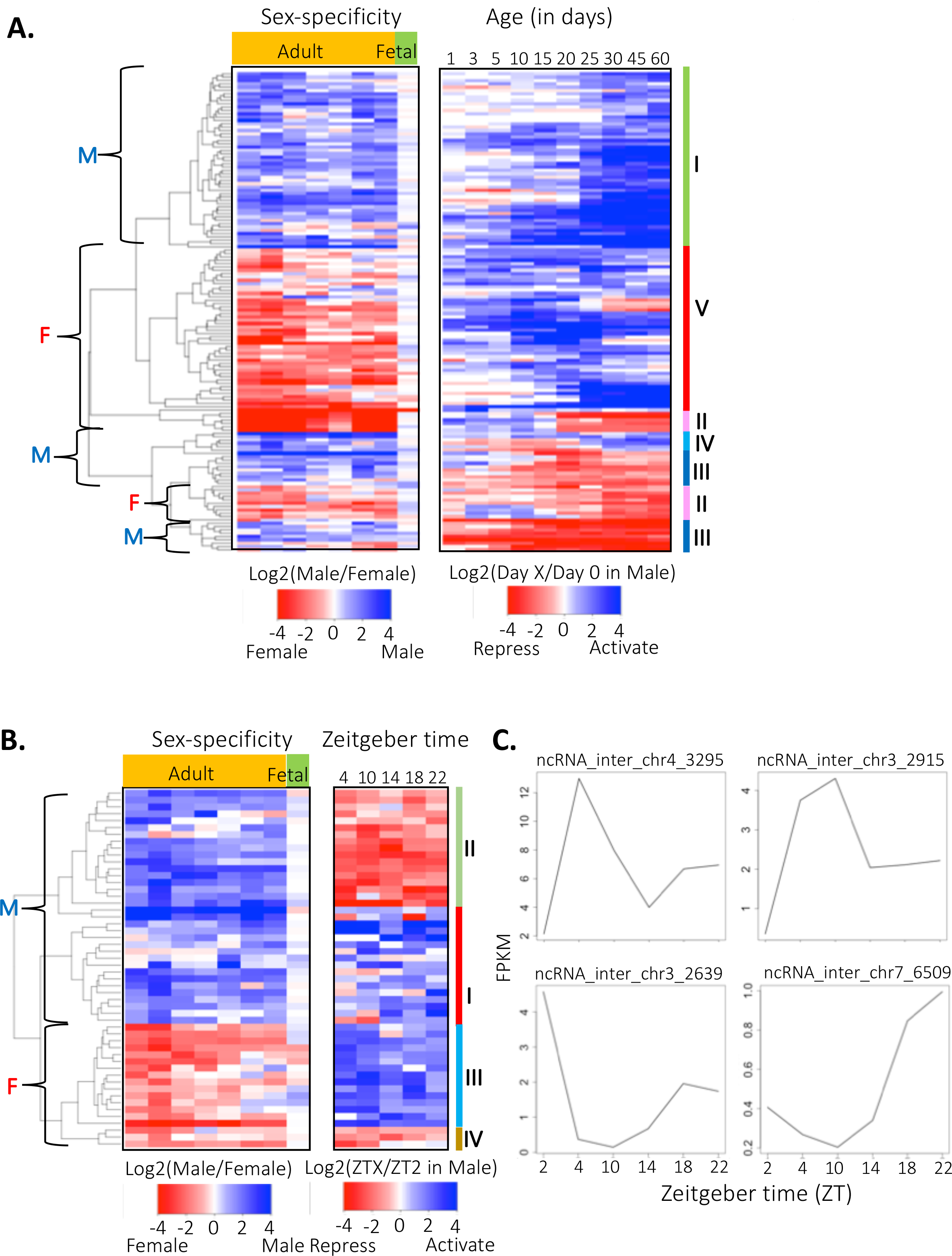
Sex-specific lncRNAs showing significant regulation during postnatal liver developmental and during a 24-hour light-dark cycle periods in male mouse liver. <u>**(A)**</u> Heatmap of 144 sex-specific lncRNAs (male/female |fold-change| > 4 at FDR < 0.05 in any of the 8 datasets assessed) that show a significant change in expression (|fold-change| > 2 and FDR < 0.05) at any time point from birth to maturity (Table S2A). The first 8 columns of the heatmap correspond to the 8 datasets to used to identify sex-specific lncRNAs; blue color indicates male-specific lncRNA expression, while red color indicates female-specific lncRNA expression. The last 10 columns track the change in expression of each lncRNA at a particular age, as compared to birth (day 0) in male liver. Blue denotes up regulation of gene expression, as compared to day 0, and red denotes down regulation. Set I designates male-specific genes that are up regulated after birth, set II are female-specific genes that are down regulated after birth, set III are male-specific genes that are down regulated after birth while maintaining their male-specific expression at adulthood, set IV are similar to set III genes, except they are up regulated around 30 days of age, and set V are female-specific genes that are up regulated by 25 days of age. See Table S2C for listings of lncRNAs in set I to V. <u>**(B)**</u> Heatmap of 52 sex-specific lncRNAs (male/female |fold-change| > 4 at FDR < 0.05 in any of the 8 datasets evaluated) that showed a significant change in expression (|fold-change| > 2 and FDR < 0.05) at one or more time points from ZT4-ZT22, as compared to ZT2 (Table S2B). The first 8 columns in the heatmap correspond to the 8 datasets used to identify sex-specific lncRNAs; blue color indicates male-specific gene expression, and red color indicates female-specific expression. The last 5 columns show the fold-change of expression at a particular ZT time, as compared to time ZT2. Blue indicates activation after ZT2; red indicates repression after ZT2. Set I and III are sex-specific lncRNAs that were up regulated at any time point after ZT2, and set II and IV are sex-specific lncRNAs that were repressed at any time point after ZT2. See Table S2D for listings of lncRNAs in set I to IV. <u>**(C)**</u> 24 h oscillating gene expression profiles are shown for four sex-specific lncRNAs, where the top figures show up regulation by ZT10, and the bottom figures show down regulation during the first 12 h.

Many organisms exhibit oscillating gene expression patterns with a period of ~24-hour (circadian rhythms). In mammals, these circadian patterns are maintained by feedback loop mechanisms orchestrated by several transcription factors [43, 68, 69]. We quantified gene expression for lncRNAs expressed in male liver, every 4 h in a 24-h period, from ZT2 (2 h after animals were exposed to light during a 12-h light and 12-h dark cycle) to ZT22 (see Methods). 241 lncRNAs, including 52 sex-specific lncRNAs, showed regulation at one or more circadian time points (|fold-change| > 2 at FDR < 0.05; Fig. 2B; Table S-42B and Table S2D). Subsets of both male-specific and female-specific lncRNAs showed higher expression at one or more time points after ZT2 (Fig. 2B, *set I and III*; Fig. 2C, *top*), and another subset showed the opposite pattern (Fig. 2B, *set II and IV;* Fig. 2C, *bottom*). We identified fewer circadian regulated female-specific lncRNAs, especially those that were down regulated after ZT2, most likely due to the exclusive use of male livers in this analysis. Overall, 157 sex-biased lncRNAs were regulated during postnatal liver development or responded to the circadian rhythm.

### Protein-coding gene clusters are enriched for biologically relevant traits

Sex-specific genes display at least two distinct phenotypes following the ablation of pituitary GH secretion by hypox, indicating there are at least two distinct classes of male-biased and two classes of female-biased genes, one subject to positive regulation and the other subject to negative regulation by GH [32, 33]. Little is known about the underlying mechanisms whereby these distinct regulatory responses to hypox are achieved. We hypothesized that sex-specific lncRNAs might play a role in this regulation. To examine this possibility, we first examined whether a strong genetic regulatory component is evident for the four distinct hypox-response classes. To accomplish this, we utilized a large panel of liver gene expression datasets available for DO mice, which have a natural high allelic variance that manifests as a variable gene expression pattern in each individual mouse. Specifically, we used WGCNA [59] to cluster sex-specific protein-coding genes based on their gene expression correlations across male DO livers, and separately, female DO livers. We identified 40 gene clusters in male livers, and 44 other gene clusters in female livers (Fig. 3 and Fig. 4; Table S3A and Table S3B). The average number of genes per cluster was comparable in the two sets of livers (25.5 genes/cluster for male livers, 23 genes/cluster for female livers). Unexpectedly, more than half of the sex-biased gene clusters were significantly enriched for either male-specific or female-specific protein coding genes (Fig. 3 and Fig. 4; *rows 4-5*), even though the analyses were performed using expression data from a single sex. Even more striking, many of the clusters were significantly enriched for one of four biologically relevant traits, defined by the change in gene expression following hypox, i.e., class 1 or class 2 male-specific genes, and class 1 or class 2 female-specific genes (Fig. 3 and Fig. 4; *rows 6-9*; Table S4A-D). These enrichments were unexpected, as the liver samples used to discover gene clusters come from DO mice, which have an intact pituitary gland, and are thus not expected to manifest the four hypox-responsive phenotypes. Further, in separate analyses, we determined that each of the 8 DO mouse founder strains is characterized by a similar distribution of sex-biased genes in each of the four hypox response classes (our unpublished results), indicating that at least a subset of GH regulatory pathways controlling sex-biased liver gene expression is present in the DO founder strains. Thus, the significant enrichment of many DO mouse sex-biased liver gene clusters for a single hypox-response class indicates that the genetic diversity of DO mice captures a key mechanistic characteristic of sex-biased gene regulation. A subset of gene regulation mechanisms related to hypox-responsiveness is presumably altered by genetic variants accumulated in DO mouse founder strains.

**Figure 3.**
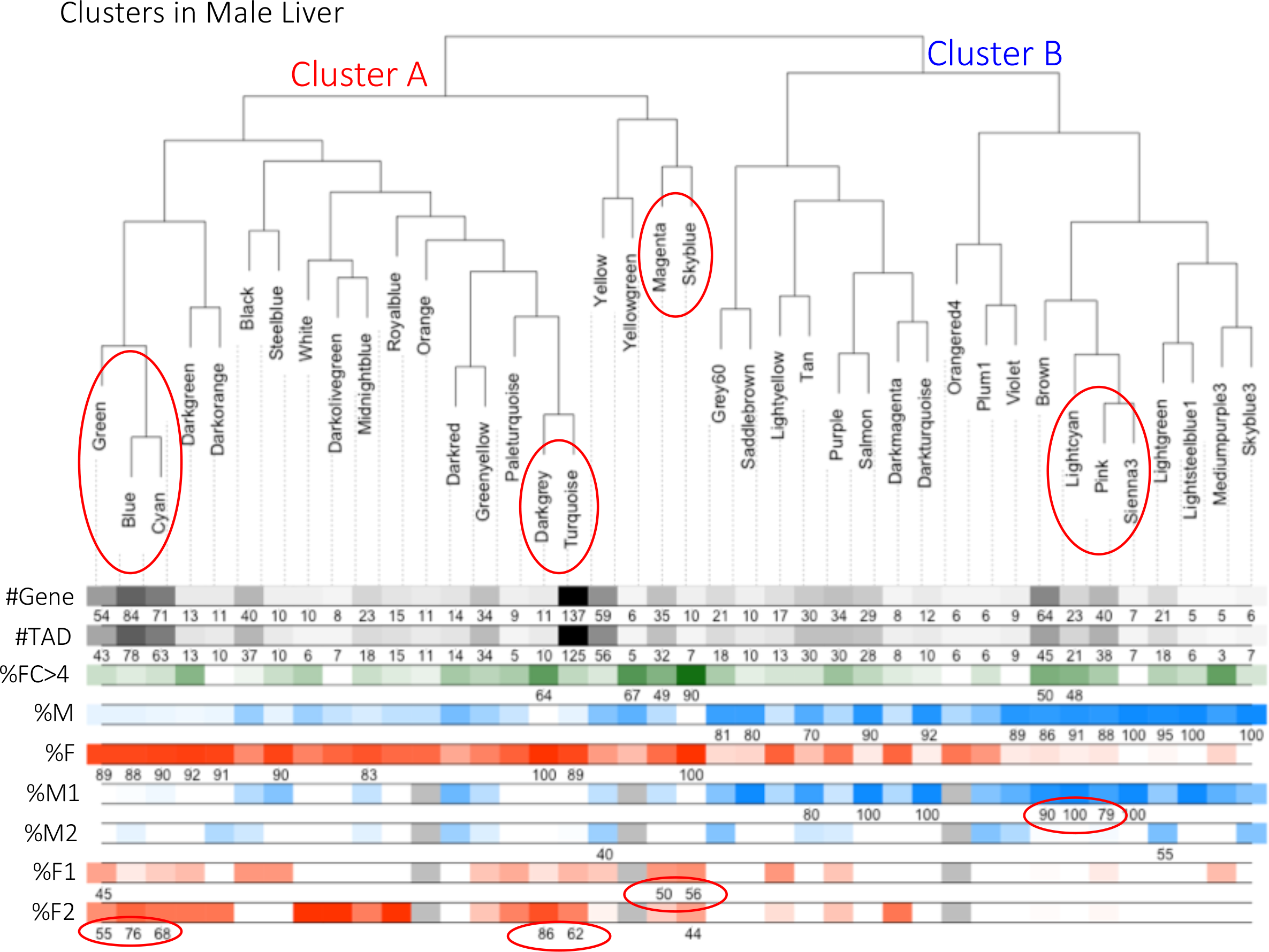
Sex-specific gene clusters (gene modules) discovered in DO male mouse liver. (*Top*) Hierarchical clustering showing the relationship between all clusters, as defined by the correlation of their first principal component. Clusters are named by *colors*, as indicated. (*Bottom*) The first row indicates the number of genes in each cluster, and the second row indicates the number of distinct TADs represented in each cluster. The next three rows show the percentage of genes in each cluster that are either strongly sex-specific (FC>4; row 3), or that are male-specific (row 4) or female-specific genes (row 5). The next four rows indicate the percentage of genes distributed in each of the four hypox-response gene classes: male-specific class 1 and class 2, and female-specific class 1 and class 2; these percentages were calculated with respect to the subset of genes in each cluster that show a significant response to hypox, consequently, the sum of percentages for the last four rows in each cluster should equal to 100%. However, for rows 3-9, the percentage values are only listed for clusters that show a significant overlap with relevant gene sets in each row (p-value < 0.05, Hypergeometric test). Ovals indicate adjacent branches whose nodes are enriched for the same hypox-response gene class. Clusters are named by *colors*, as indicated; in several cases the cluster name (i.e., color) assigned to a cluster, discovered in DO male mouse liver is the same name as that assigned to one of the clusters discovered in DO female mouse liver and shown in Fig. 4; however, there is no relationship between such clusters. Thus, male DO mouse liver cluster Cyan, with 71 genes, displayed here, is not related to female DO mouse liver cluster Cyan, with 20 genes, shown in Fig. 4. Cluster A consists of protein-coding gene modules enriched for female-specific genes while cluster B consists of protein-coding gene modules enriched for male-specific genes.

**Figure 4.**
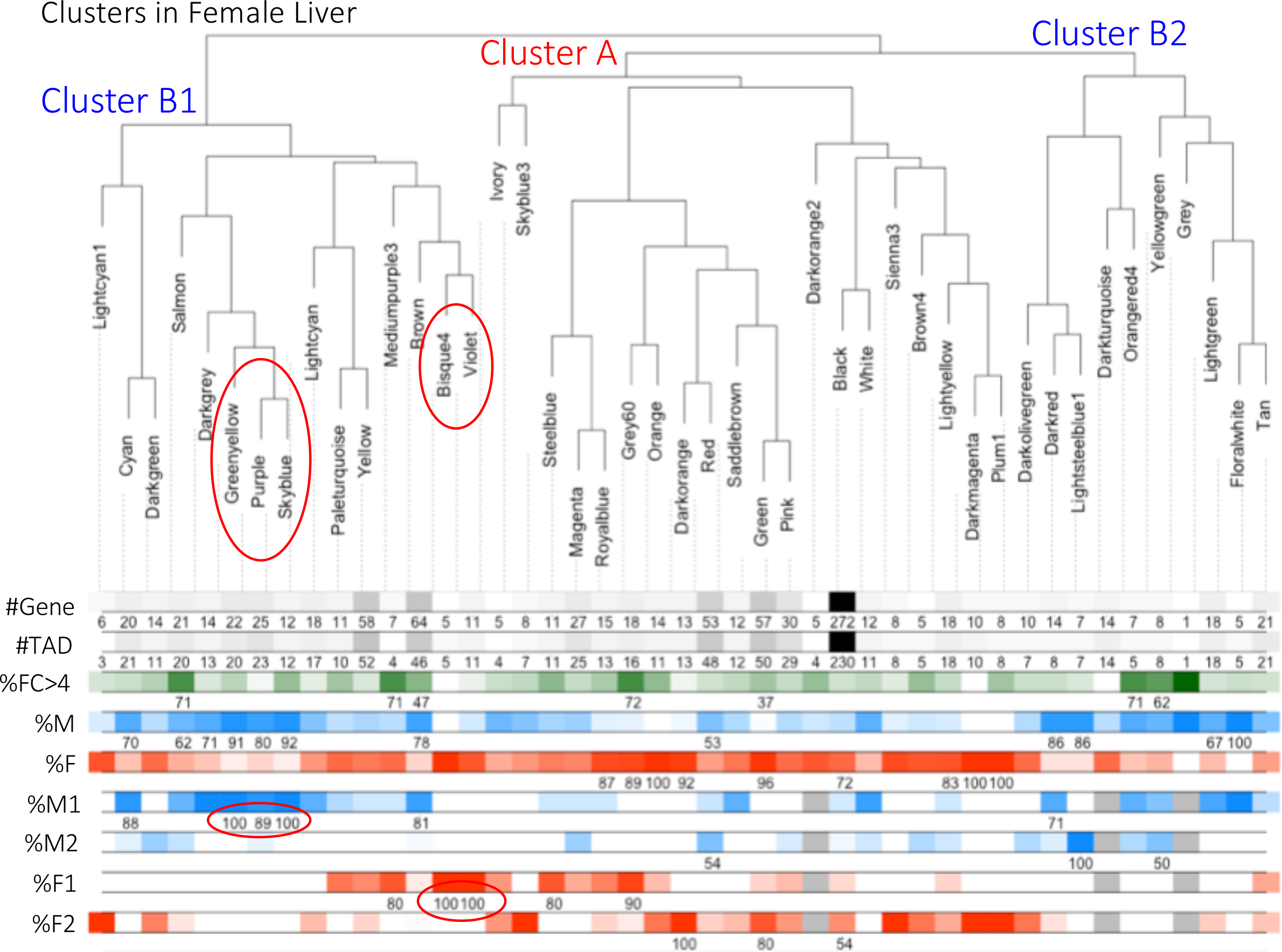
Sex-specific gene clusters discovered in DO female mouse liver. Characteristics of the gene clusters shown are displayed as described in the legend to Fig. 3. Cluster A consists of protein-coding gene modules enriched for female-specific genes while cluster B1 and B2 consist of protein-coding gene modules enriched for male-specific genes.

Overall, in DO male livers, 13 gene clusters encompassing 240 male-specific protein-coding genes were significantly enriched (p < 0.05, Hypergeometric test) for male-specific genes, and an additional 10 clusters encompassing 379 female-specific protein-coding genes were significantly enriched for female-specific genes (Table 1). Furthermore, a subset was significantly enriched for strongly sex-specific genes (|fold-change| > 4; Fig. 3 and Fig. 4; *row 3*). Significant enrichments were also found in DO male liver for the four hypox-response gene classes (Table 1): 7 gene clusters (encompassing 85 genes) were enriched for male class 1 genes, 2 gene clusters (encompassing 12 genes) were enriched for male class 2 genes, 3 clusters (encompassing 21 genes) were enriched for female class 1 genes, and 6 clusters (encompassing 89 genes) were enriched for female class 2 genes. 14 of 16 clusters that are enriched for one of the four classes of hypox-responsive genes were enriched for sex-specific genes. Protein-coding gene clusters with similar patterns of enrichment for sex-biased genes, and for hypox-response gene classes, were discovered in DO female liver samples, as is summarized in Fig. 4 and Table 1. The identified clusters do not consist of genes from the same genomic regions, as a majority of genes in every cluster are from different topologically associated domains (TADs) [70] (Fig. 3 and Fig. 4; *row 2*), indicating the potential for this approach to identify *trans* regulation.

**Table 1.** Enrichments of gene modules discovered in male, and separately, in female liver for sex-specific and hypox-responsive genes. #Cluster columns indicate the number of cluster that are enriched for each respective gene set. The #Gene columns indicate the number of genes belonging to each respective gene set in these enriched cluster. The common genes row indicates the number of genes that overlap in male and female liver in each column.

| Samples | Enriched for Male-specific Genes |  | Enriched for Female-specific Genes |  | Enriched for Male Class I Genes |  | Enriched for Male Class II Genes |  | Enriched for Female Class I Genes |  | Enriched for Female Class II Genes |  |
| --- | --- | --- | --- | --- | --- | --- | --- | --- | --- | --- | --- | --- |
|  | #Cluster | #Gene | #Cluster | #Gene | #Cluster | #Gene | #Cluster | #Gene | #Cluster | #Gene | #Cluster | #Gene |
| Male | 13 | 240 | 10 | 379 | 7 | 85 | 2 | 12 | 3 | 21 | 6 | 89 |
| Female | 12 | 201 | 9 | 339 | 6 | 68 | 3 | 11 | 5 | 23 | 3 | 82 |
| Common genes | 143 |  | 265 |  | 56 |  | 6 |  | 7 |  | 68 |  |

Although overall cluster statistics and enrichments were similar between male and female liver (Table 1), ~82% (33 of 40) of the gene clusters discovered in male liver did not have any counterpart in the female liver clusters; only 7 of the 40 DO male liver clusters overlap (defined as >50% genes in common) with any cluster discovered in female liver (Fig. S4). Five of the 7 overlapping clusters (circled in Fig. 5) were enriched for either male-specific or female-specific genes, but not for highly sex-specific genes. An earlier study, using another mouse strain, found a higher overlap (53%) between gene clusters in the two sexes, which may be due to those analyses clustering all varying genes, as opposed to only sex-specific genes. The degree of gene expression correlation within a cluster varied among cluster members (Fig. S5), with some but not other genes showing strong correlations with many genes in the cluster. Thus, there is an inherent structure of gene regulation for each cluster, where genes with many connections to other genes may play an important role in regulation.

**Figure 5.**
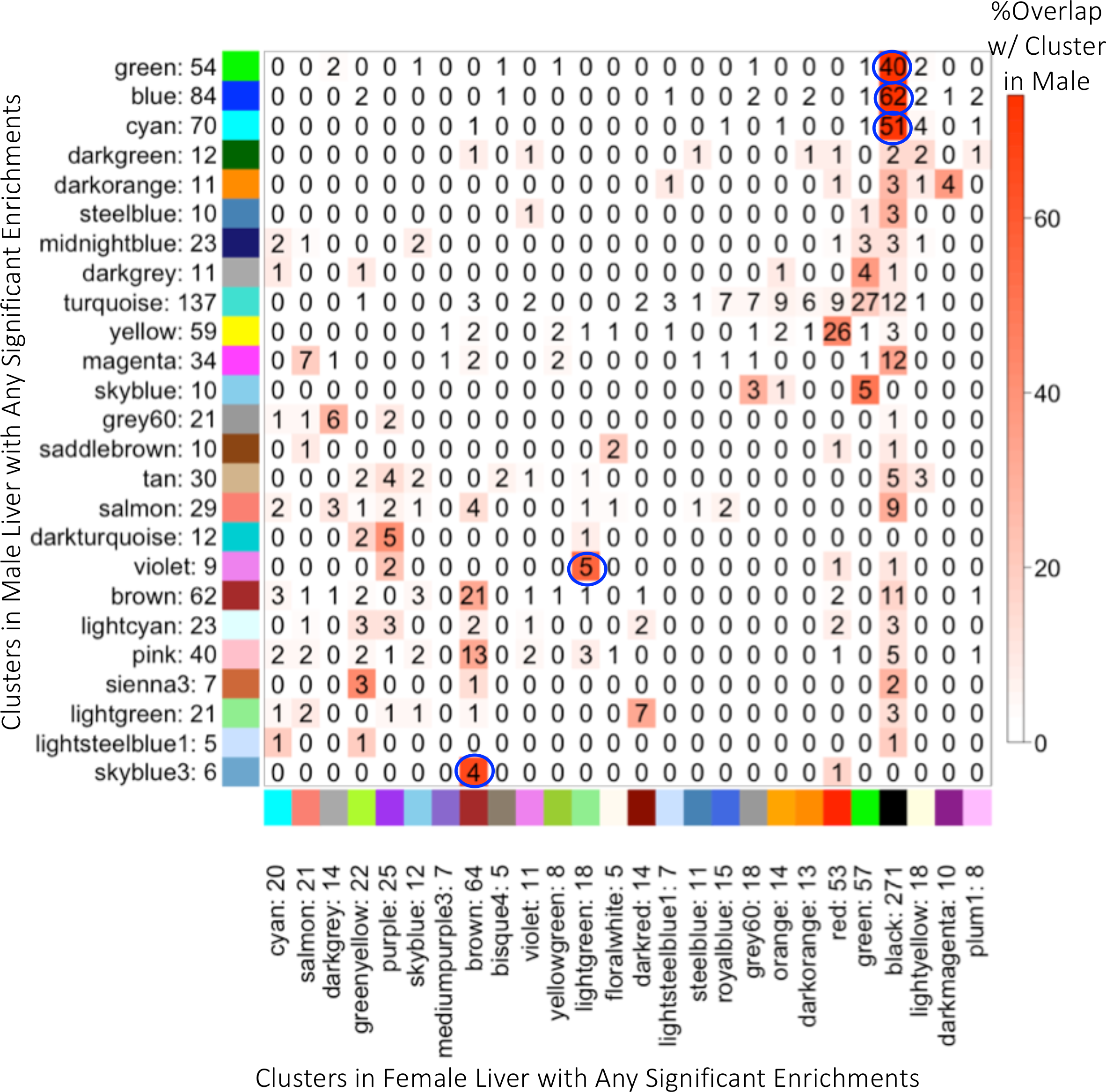
Overlap between clusters discovered in male *vs*. female liver for clusters that are significantly enriched for sex-specific genes and/or hypox-response gene classes. The number shown next to each cluster name (designated as a *color*; see legend to Fig. 3) represents the number of genes in that cluster. Clusters discovered in DO male liver are shown along the Y-axis; those discovered in DO female liver are shown along the X-axis. The matrix shows the number of genes common to the intersecting gene clusters discovered in the DO livers from each sex. Shades of red (bar at *right*) represent the percentage of overlap with respect to the cluster discovered in male liver. Circles denote cluster pairs with strong gene overlap, defined as >50% of gene members in a specific cluster in DO male livers (row) being present in a specific cluster in DO female livers (column).

### Sex-biased lncRNAs negatively correlate with sex-opposite and inverse hypox class-enriched protein-coding gene clusters

Next, we considered whether any of the 168 multi-exonic, intergenic sex-specific lncRNAs identified above might serve as a regulator of any of the sex-specific protein-coding gene clusters (gene modules), as indicated by gene expression correlations. Significant correlations (FDR < 0.001) were seen between sex-biased protein coding gene modules and 73 sex-specific lncRNAs, with 40 of these lncRNAs showing correlations in DO male liver (Table S5A) and 52 showing correlations in DO female liver (Table S5B). Most of these were positive correlations, which could indicate co-expression/co-regulation between the lncRNA and the protein-coding gene module. We therefore focused on negative regulators, because of the difficulty of distinguishing a lncRNA that serves as a positive regulator of a gene module from a lncRNA that is simply a co-expressed gene. We identified male class 1 lncRNAs whose expression across DO mouse livers showed a significant negative correlation with protein-coding gene clusters enriched for female class 2 genes in male liver, as well as female class 1 lncRNAs that were negatively correlated with male class 2-enriched protein-coding gene modules in female liver, and correspondingly for class 2 sex-specific lncRNAs (Fig. 6). A negative correlation of gene expression profiles between inverse hypox classes of the opposite sex is consistent with the models shown in Fig. 6, and would enable a lncRNA to negatively regulate protein-coding genes by one of several established mechanisms [24, 71, 72]. 16 sex-biased multi-exonic lncRNAs showed significant (FDR < 0.001) negative correlations with gene clusters enriched (P < 0.01) with sex-opposite protein coding genes of the inverse hypox class (Table 2 and Table S8), consistent with the proposed negative regulatory role for these lncRNAs (c.f. Fig. 6). In some cases, the correlations involve male-specific lncRNAs whose expression in female DO liver negatively correlated with clusters enriched for highly female-specific protein-coding genes (e.g., ncRNA_inter_chr10_9000) or female-specific lncRNAs whose expression in male DO liver negatively correlated with clusters enriched for highly male-specific protein-coding genes (e.g., ncRNA_inter_chr2_1430) (Table 2). In these cases, low levels of the male-specific lncRNA would need to be maintained in female liver in order for the inverse hypox class, female-specific gene module to maintain its high expression in female liver, and correspondingly for female-specific lncRNAs and inverse hypox class, male-specific gene modules in male liver. Overall, the negatively correlated sex-biased gene clusters shown in Table 2 encompass a substantial fraction (327 of 1,019) of sex-biased protein coding genes included in our clusters (182 male-biased genes and 145 female-biased genes; 75 male class 1, 11 male class 2, 22 female class 1 and 35 female class 2 genes). Example of individual clusters are shown in Fig. S5.

**Figure 6.**
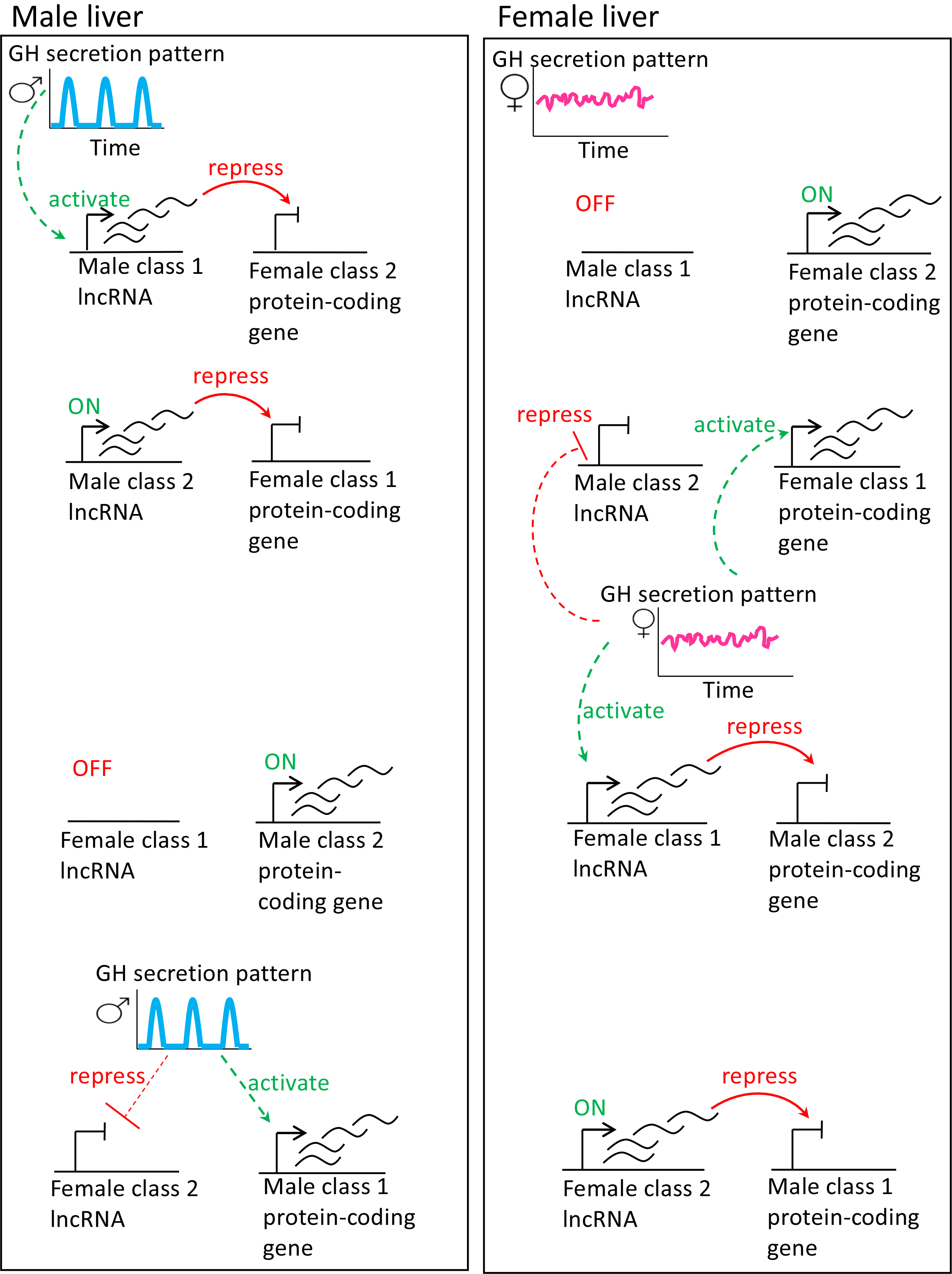
Schematic diagram of the eight proposed scenarios for negative regulatory role of sex-specific lncRNAs. (*Left*) Gene regulatory mechanisms in male liver: the male, pulsatile plasma GH pattern activates male class 1 lncRNAs in male liver, which are proposed to repress female-class 2 protein-coding genes in male liver (*first row*). Similarly, male-class 2 lncRNAs, which are constitutively expressed in male liver, are proposed to repress female class 1 protein-coding genes in male liver (*second row*). Female class 1 lncRNAs are constitutively off in male liver, and therefore cannot repress male-class 2 protein-coding genes (*third row*). Female class 2 lncRNAs, whose expression is repressed by the male plasma GH pattern in male liver, therefore cannot repress male class 1 protein-coding genes (*fourth row*). (*Bottom*) Gene regulatory mechanisms in female liver: in the absence of pulsatile GH, male class 1 lncRNAs are constitutively off in female liver, and thus cannot repress female-class 2 protein-coding genes (*top row*). Male-class 2 lncRNAs are repressed by the female plasma GH pattern, and cannot therefore repress female class 1 protein-coding genes (*second row*). The female, persistent plasma GH pattern activates female class 1 lncRNAs, which are proposed to repress male-class 2 protein-coding genes (*third row*). Female-class 2 lncRNAs are constitutively on in female liver, and can therefore repress male class 1 protein-coding genes (*fourth row*). The inverse regulatory relationship between class 1 sex-specific lncRNAs and class 2 sex-specific protein coding genes shown here is consistent with the effects of hypox on both classes of sex-specific genes. For example, hypox of male mice, which abolishes the GH pulse regulatory circuit diagrammed on the *left*, down regulates class 1 male lncRNAs and up regulates (de-represses) female class 2 protein coding genes in male liver. That de-repression is proposed to involve the loss of the male class 1 lncRNAs required for female class 2 protein-coding gene repression. Hypox of female mice, which abolishes the persistent GH regulatory circuit diagrammed on the *right*, down regulates class 1 female lncRNAs and up regulates (de-represses) male class 2 protein coding genes in female liver.

**Table 2.** List of sex-specific IncRNAs that are negatively correlated (FDR < 0.001) with protein-coding genes enriched (P < 0.01) for the inverse hypox response classes

| Liver data set | lncRNA | Hypox response class of the lncRNA | Pearson correlation coefficient | Significance of correlation (FDR) | Sex-specific protein-coding gene module | Hypox response class that shows enrichment in the module | Significance of the hypox class enrichment for each module | # Genes in the module |
| --- | --- | --- | --- | --- | --- | --- | --- | --- |
| Female | ncRNA_inter_chr7_6509 | Male I | -0.59 | 1.05E-09 | darkorange | Female II | 2.47E-03 | 13 |
| Female | ncRNA_inter_chr1_630 | Male I | -0.45 | 3.82E-05 | green | Female II | 3.17E-06 | 57 |
| Female | ncRNA_inter_chr10_9313 | Male I | -0.42 | 1.34E-04 | darkorange | Female II | 2.47E-03 | 13 |
| Female | ncRNA_inter_chr6_5316 | Male I | -0.47 | 1.01E-05 | green | Female II | 3.17E-06 | 57 |
| Male | ncRNA_inter_chr8_7423 | Male I | -0.45 | 1.08E-04 | darkgrey | Female II | 8.01E-04 | 11 |
| Female |  |  | -0.44 | 5.70E-05 | green | Female II | 3.17E-06 | 57 |
| Female | ncRNA_inter_chr10_9000 | Male II | -0.60 | 3.35E-10 | grey60 | Female I | 1.32E-06 | 18 |
| Female | ncRNA_inter_chr16_13349 | Male II | -0.40 | 3.14E-04 | steelblue | Female I | 6.42E-03 | 11 |
| Female | ncRNA_inter_chr8_6946 | Female I | -0.53 | 1.73E-07 | red | Male II | 2.17E-03 | 53 |
| Female | ncRNA_inter_chr13_11437 | Female I | -0.49 | 1.93E-06 |  |  |  |  |
| Female | ncRNA_inter_chr16_13428 | Female I | -0.41 | 2.90E-04 |  |  |  |  |
| Male | ncRNA_inter_chr1_307 | Female II | -0.40 | 8.55E-04 | lightcyan | Male I | 1.91E-04 | 23 |
| Female | ncRNA_inter_chr10_9222 | Female II | -0.47 | 1.01E-05 | skyblue | Male I | 1.68E-03 | 12 |
| Female |  |  | -0.39 | 6.14E-04 | purple | Male I | 8.23E-03 | 25 |
| Male | ncRNA_inter_chr16_13510 | Female II | -0.44 | 1.44E-04 | pink | Male I | 3.57E-05 | 40 |
| Male | ncRNA_inter_chr2_1430 | Female II | -0.51 | 4.04E-06 |  |  |  |  |
| Male |  |  | -0.48 | 1.18E-05 | lightcyan | Male I | 1.91E-04 | 23 |
| Male |  |  | -0.43 | 1.82E-04 | sienna3 | Male I | 4.98E-04 | 7 |
| Male |  |  | -0.41 | 4.53E-04 | brown | Male I | 9.58E-11 | 64 |
| Male | ncRNA_inter_chr2_1451 | Female II | -0.45 | 1.08E-04 | lightcyan | Male I | 1.91E-04 | 23 |
| Male |  |  | -0.42 | 3.18E-04 | pink | Male I | 3.57E-05 | 40 |
| Female | ncRNA_inter_chr3_2937 | Female II | -0.58 | 2.63E-09 | purple | Male I | 8.23E-03 | 25 |
| Female |  |  | -0.54 | 1.27E-07 | greenyellow | Male I | 1.96E-06 | 22 |
| Female |  |  | -0.41 | 2.62E-04 | skyblue | Male I | 1.68E-03 | 12 |

Negative correlations between sex-opposite lncRNAs and inverse hypox class-enriched protein-coding gene clusters (Table 2) were also seen when we included more weakly correlated (FDR < 0.05) lncRNAs, particularly in male liver (Fig. 7, *right* panels). Fig. 7A plots the hypox class distributions for 94 sex-specific lncRNAs that show positive correlations (*left*) or negative correlations (*right*) with 16 protein-coding gene modules that are enriched for one of the four hypox-response classes. As expected, more male class 1 lncRNAs are positively correlated with gene modules that show strong enrichment for male class 1 protein-coding genes, as compared to those that do not (Fig. 7A *left*; first eight bars *vs* the other bars), or as compared to the overall distribution for the 142 sex-specific lncRNAs that are varying in male liver (MAD > 0; Fig. 7A *All;* Table S2E). A similar pattern was seen in female DO livers (Fig. 7B *left*; first six bars *vs* the others, and Fig. 7B *left* vs *All*; Table S2F). Negative correlations between inverse hypox-response classes were also seen, with more female class 2 lncRNAs being negatively correlated with protein coding gene modules enriched for male class 1 protein-coding genes, as compared to those that are not (Fig. 7A *right*; first seven bars *vs* the others) or as compared to the overall distribution for all sex-specific lncRNAs that are varying in male liver (Fig. 7A *right* vs *All*; Table S2E). A similar, albeit less prominent, negative correlation between female class 1 lncRNAs and protein-coding gene modules enriched for male class 2 genes was observed (Fig. 7B *right*; first two bars *vs* the others). Taken together, these findings support a model whereby class 1 sex-biased lncRNAs negatively regulate protein-coding modules that are enriched for class 2 sex-biased genes of the opposite sex-specificity, and vice versa.

**Figure 7.**
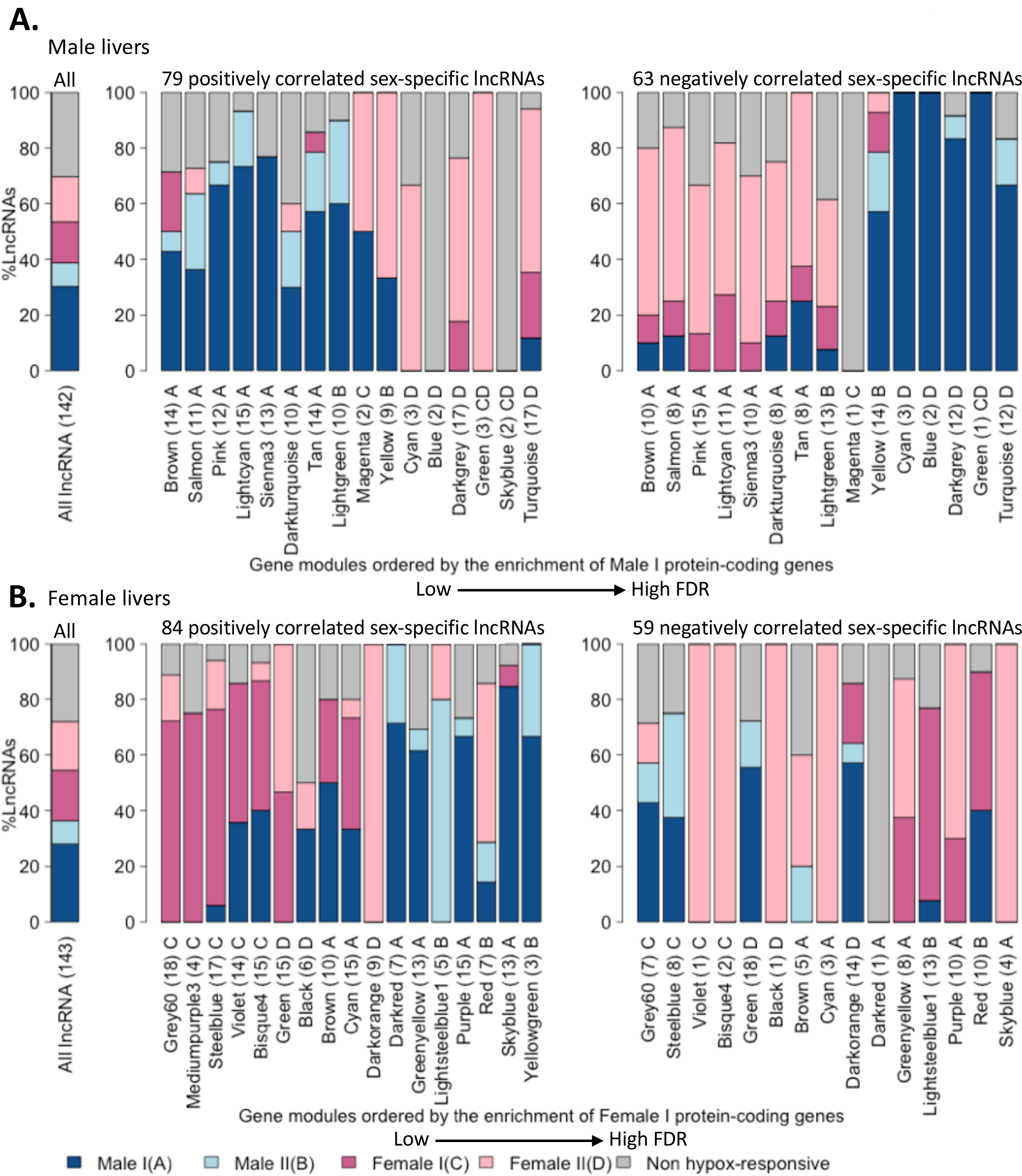
Distribution of hypox-response gene classes for intergenic, sex-specific lncRNAs whose expression in DO mouse liver is eitherpositively or negatively correlated with protein-coding gene modules that are enriched for one the four hypox-response classes. <u>**(A)**</u> Shown is the distribution of hypox-response classes for the 142 multi-exonic, intergenic sex-specific lncRNAs that are varying (MAD > 0) in male liver (first bar, marked *All*), 79 sex-specific lncRNAs whose expression is positively correlated with any protein-coding gene module enriched for hypox-responsive genes in male liver (*left*) and 63 sex-specific lncRNAs that are negatively correlated with any protein-coding gene module enriched for hypox-responsive genes in male liver (*right*). Gene modules in the *left* and *right* panels are both ordered by their enrichment for male class 1 protein-coding genes from left to right (low FDR to high FDR). More male class 1 lncRNAs are seen to be positively correlated with protein-coding genes that are strongly enriched for male class 1 genes, whereas more female class 2 lncRNAs are negatively correlated with protein coding gene clusters enriched for male class 1 genes. <u>**(B)**</u> Shown is the distribution of hypox-response classes for the 143 multi-exonic, intergenic sex-specific lncRNAs that are varying (MAD > 0) in female liver (first bar, marked *All*), 84 sex-specific lncRNAs that are positively correlated with any protein-coding gene modules enriched for hypox-responsive genes in female liver (*left*), and 59 sex-specific lncRNAs that are negatively correlated with any protein-coding gene modules enriched for hypox-responsive genes in female liver (right). Gene modules in the *left* and *right* panels are both ordered by their enrichment for female class 1 protein-coding genes from left to right (low FDR to high FDR). The number in parenthesis after each gene cluster name indicates the total number of lncRNAs whose hypox class distributions are shown in each bar graph. More female class 1 lncRNAs positively correlate with protein-coding genes that are strongly enriched for female class 1 genes, and more, albeit to a lesser extent, male class 2 lncRNAs negatively correlate with protein coding gene clusters enriched for female class 1 genes.

## Discussion

Many genes in mammalian liver are differentially expressed between males and females [1–3, 31], including lncRNA genes [22], which are thought to contribute to liver sex differences by establishing or maintaining the widespread sex-differences in chromatin accessibility and chromatin states that are closely linked to sex differences in liver transcription factor binding and chromatin states [16, 20, 21, 73]. Here, we expanded the repertoire of liver-expressed ncRNAs to include more than 15,500 lncRNAs, including many novel genes, based on liver gene expression data collected under 30 distinct biological conditions. A subset of these lncRNAs showed strong sex-biased expression, as well as regulation during liver development or in response to circadian rhythm. Further, we identified sex-specific lncRNAs whose expression across DO mouse livers shows a significant negative correlation with protein-coding gene modules enriched for the genes of the opposite sex-bias and inverse hypox-response class, consistent with the proposal that these lncRNAs play an important negative regulatory role in controlling sex-biased gene expression. The broad characteristics of sex-biased gene expression revealed by this study include an indication that multiple, distinct gene regulatory mechanisms likely contribute to the four classes of hypox-responsive genes. These findings illustrate the usefulness of gene modules for generating hypotheses for lncRNA function, and reveal underlying characteristics governing sex-biased gene regulation.

### Regulation of lncRNA gene expression in male liver

We examined the expression of sex-biased lncRNAs during postnatal male liver development, where a majority of changes relevant to the emergence of liver sex-biased gene expression have been found to take place [30]. We found that a majority of male-specific lncRNAs were up regulated postnatally, as early as the first week of life. Further, we identified two patterns of up regulation for female-specific lncRNAs: sustained or dynamic up regulation after birth, with the latter pattern showing either down regulation or reduced up regulation of gene expression around 4 wk of age. These sex-specific lncRNAs may contribute to the acquisition of sex-biased gene expression at puberty and in early adulthood. We also evaluated the regulation of sex-specific lncRNAs by circadian rhythms in male liver, which gives rise to a diurnal rhythm affecting the expression of a large number of liver-expressed protein coding genes [43, 44]. Sex-biased lncRNAs showing significant gene expression changes in response to the alternating light and dark cycle in male liver could contribute to the regulation sex-specific protein-coding genes with a matching circadian rhythm, which characterizes a subset of liver sex-biased genes [43].

### Complex regulation characterizes the four hypox-response gene classes

We identified protein-coding gene modules in male liver, and separately, in female liver, of which almost half of the identified clusters are enriched for either male-specific or female-specific genes. Hierarchical clustering of these gene modules showed that, both in male and in female DO mouse liver, almost all of the modules that are enriched for male-specific genes segregated into their own branch, separated from the gene modules that are enriched for female-specific genes (Fig. 3, Fig. 4). This indicates that genetic variation in the population has a major impact on sex-biased gene expression, which in the case of DO mice is easy to discern given the genetic variability in this mouse population. One mechanism for genetic factors to exert their regulatory effect in a sex-biased manner is through genetic variants that are within GH-dependent open chromatin regions (DHS) or GH-regulated TF binding sites. GH has been shown to impart sex differences in liver through its regulation of several GH-dependent transcription factors [1, 14, 16, 17, 19, 74]. On the other hand, gene modules that were enriched for one of the four hypox-response gene classes were found in multiple, separate branches, and mixture of both class 1 and class 2 genes can be found within the same branch (Fig. 3, Fig. 4). Each co-regulated gene cluster enriched for one of the four hypox-response classes is thus likely to be distinctly regulated, which in turn shapes their gene expression pattern differently across DO mice, as compared to other gene clusters enriched for the same hypox-response gene class. This suggests that multiple, distinct regulatory mechanisms are at play in shaping each of the four major classes of hypox-responsive genes, highlighting the complexity of their regulation. Further characterization of the identified gene clusters may include discovery, for each gene cluster, of enrichments for TF binding sites, developmental time points, or circadian groups, all of which may help delineate molecular mechanisms characterizing the four hypox-response classes. Only 7 of the protein-coding gene clusters discovered in male and female liver showed substantial overlap with each other. These 7 gene clusters encompass 223 of the 1,033 sex-specific genes considered (numbers based on clusters discovered in male liver), indicating that a majority of gene regulation governing sex-biased expression involves sex-differential mechanisms.

### Sex-biased lncRNAs are proposed to negatively regulate select protein-coding gene modules

73 multi-exonic, intergenic sex-specific lncRNAs showed a highly significant correlation (FDR < 0.001) with at least one sex-specific protein-coding gene module. Further, 16 of these sex-specific lncRNAs showed significant negative correlation with a protein-coding gene module(s) enriched for genes with the opposite sex-bias and inverse hypox-response class, supporting the hypothesis that these lncRNAs negatively regulate their inversely correlated protein-coding gene modules. The eight scenarios, where negative gene expression correlations between inverse hypox classes indicate a negative regulatory role (Fig. 6), can be broadly divided into two groups: in one group, high expression of a sex-specific lncRNA is needed to repress protein-coding genes of the opposite sex-bias, e.g., male class 1 lncRNAs and female class 2 gene modules in male liver; and in the second group, low expression of a sex-specific lncRNA is needed to prevent repression of protein-coding genes of the opposite sex bias, e.g., male class 1 lncRNAs and female class 2 gene modules in female liver. These strong negativ correlations between inverse hypox classes were not expected to be seen in DO mice, which have an intact pituitary gland and presumably maintain sex-specific plasma GH patterns are. Genetic variants that have accumulated in individual DO mice, an outbred population derived from 8 inbred founder mouse stains [45, 75], must therefore alter a subset of genomic regulatory regions that have a substantial impact on the hypox-response phenotypes. This conclusion is supported by our recent finding that DO mouse founder strain genetic variants are associated with gene expression changes in select correlated lncRNA—protein coding gene module pairs (our unpublished studies). One example is the darkgrey cluster in male liver, a female class 2-enriched gene module, which negatively correlates with ncRNA_inter_chr8_7423, a male class 1 lncRNA (Table 2). eQTLs identified for both this lncRNA and for 7 of its 11 gene cluster members point to the genotype of the PWK/PhJ strain as being associated with the largest changes in expression of the protein-coding genes (*Hao2*, *Cyp2a22*, *Sult1e1*, *Sybu*, *Tm4sf4*, *Setbp1* and *Dcbld1*; Table S4A). Further, eQTLs for 3 of the other 4 gene cluster members identified the PWK/PhJ strain as the strain whose genetic variants make the second to fourth largest contribution to the observed changes in each of the 3 protein-coding gene expression (*Cyp2b9*, *Cyp2a4* and *Cep112*). Many of these eQTLs map to different chromosomes, consistent with this male-specific lncRNA being involved in *trans* regulation.

In conclusion, we expanded our earlier characterization of liver-expressed lcnRNAs by using gene expression data obtained under 30 distinct biological conditions, and identified many lncRNAs that show strong sex-biased gene expression. A subset of these sex-specific lncRNAs is dynamically regulated during postnatal liver development in male mice, and an overlapping subset shows significant oscillating gene expression patterns over a 24-hour period. We further identified sex-specific protein-coding gene clusters in male liver, and separately, in female liver using a mouse population with rich genetic diversity (DO mouse), many of which showed unexpected enrichment for one of the four hypox-response classes. Sex-specific lncRNAs that are significantly correlated in expression with at least one protein-coding gene cluster were identified. Strikingly, a subset comprised of 16 sex-specific lncRNAs showed a significant negative correlation with protein-coding gene modules that were enriched for genes of the opposite sex-bias and inverse hypox classes, suggesting a negative regulatory role for these lncRNAs. These findings elucidate the sex-bias liver transcriptome, and illustrate the utility of gene modules to reveal broad characteristics of sex-bias gene regulation and generate hypothesis for regulatory roles for sex-bias lncRNAs.

## Acknowledgements

The authors thank Christine Nykyforchyn of the Waxman lab for her analysis of GH-responsive lncRNAs using the ribosomal-depleted RNA-Seq samples and her initial analysis of the gene expression correlations in DO mouse liver.

## Conflict of Interest

The authors declare that they have no competing interests.

## Supplemental Figure legends

**Figure S1.**
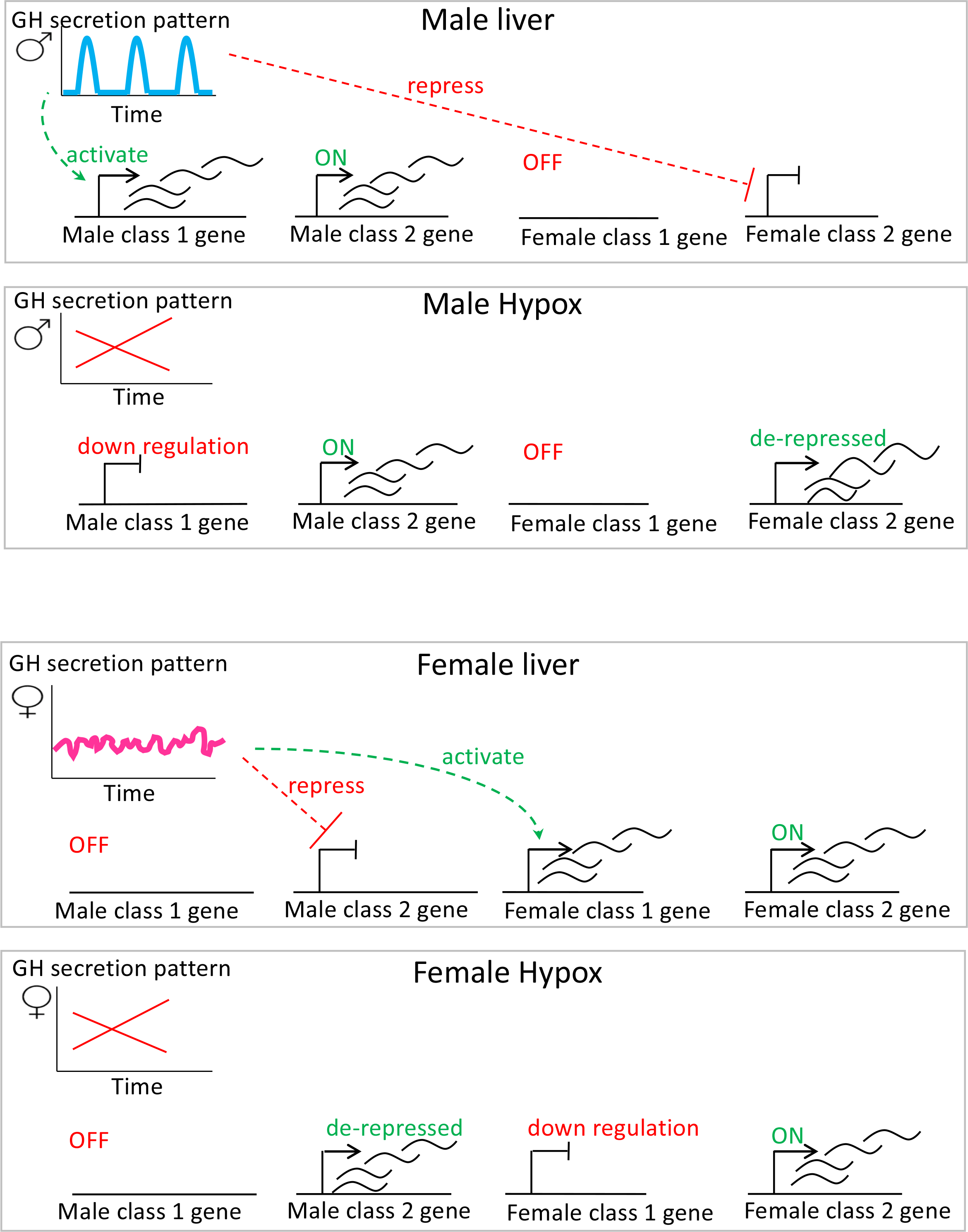
Four hypox-response classes. Genes in male hypox class 1 and female hypox class 1 are activated by the GH secretion pattern of the sex where they are more highly expressed, whereas, male hypox class 2 and female hypox class 2 genes are repressed by the GH secretion pattern of the sex where they are less highly expressed. Consequently, following hypox, class 1 genes are down regulated and class 2 genes are de-repressed (up regulated).

**Figure S2.**
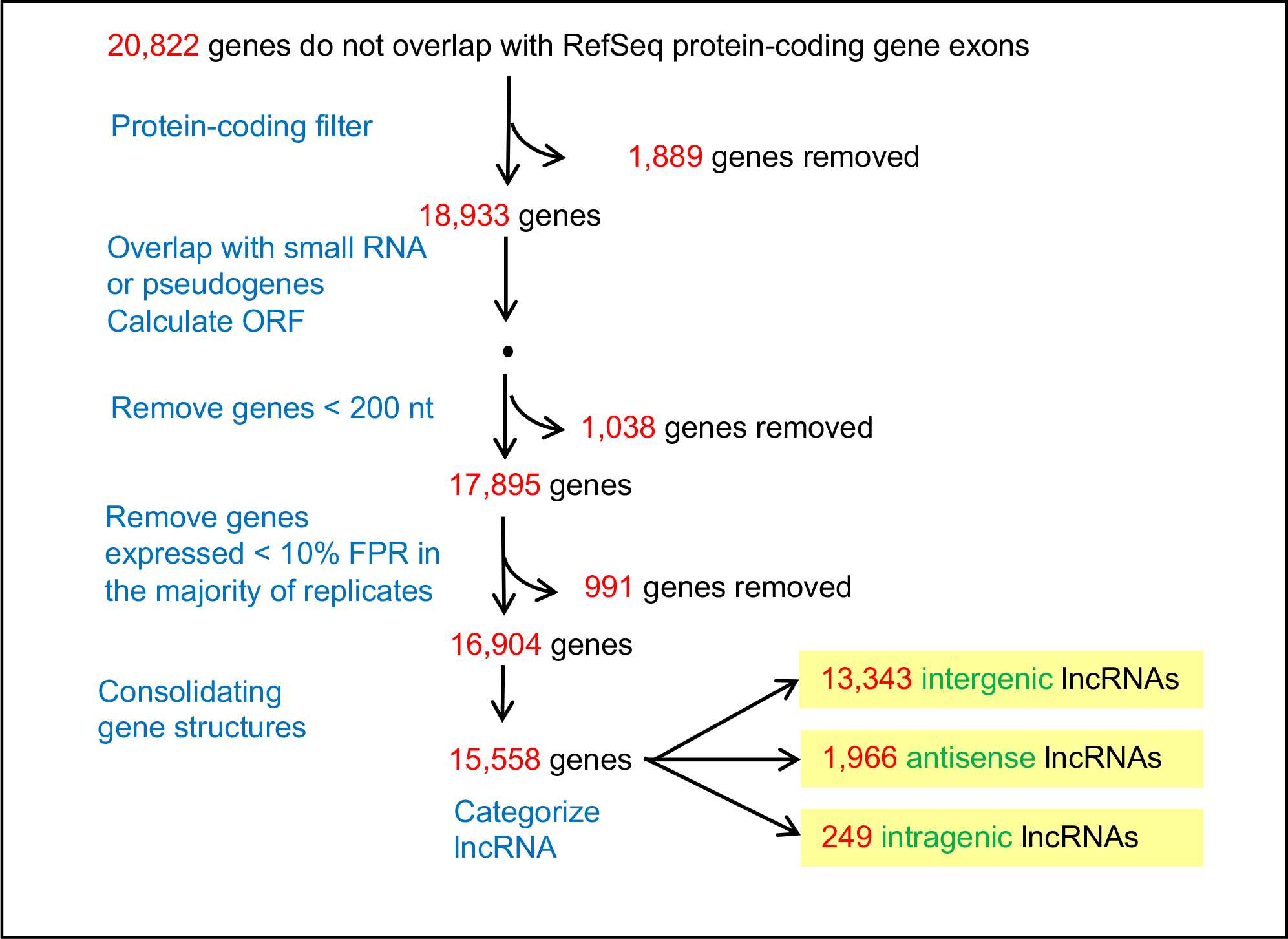
Pipeline to discover liver-expressed lncRNAs.

**Figure S3.**
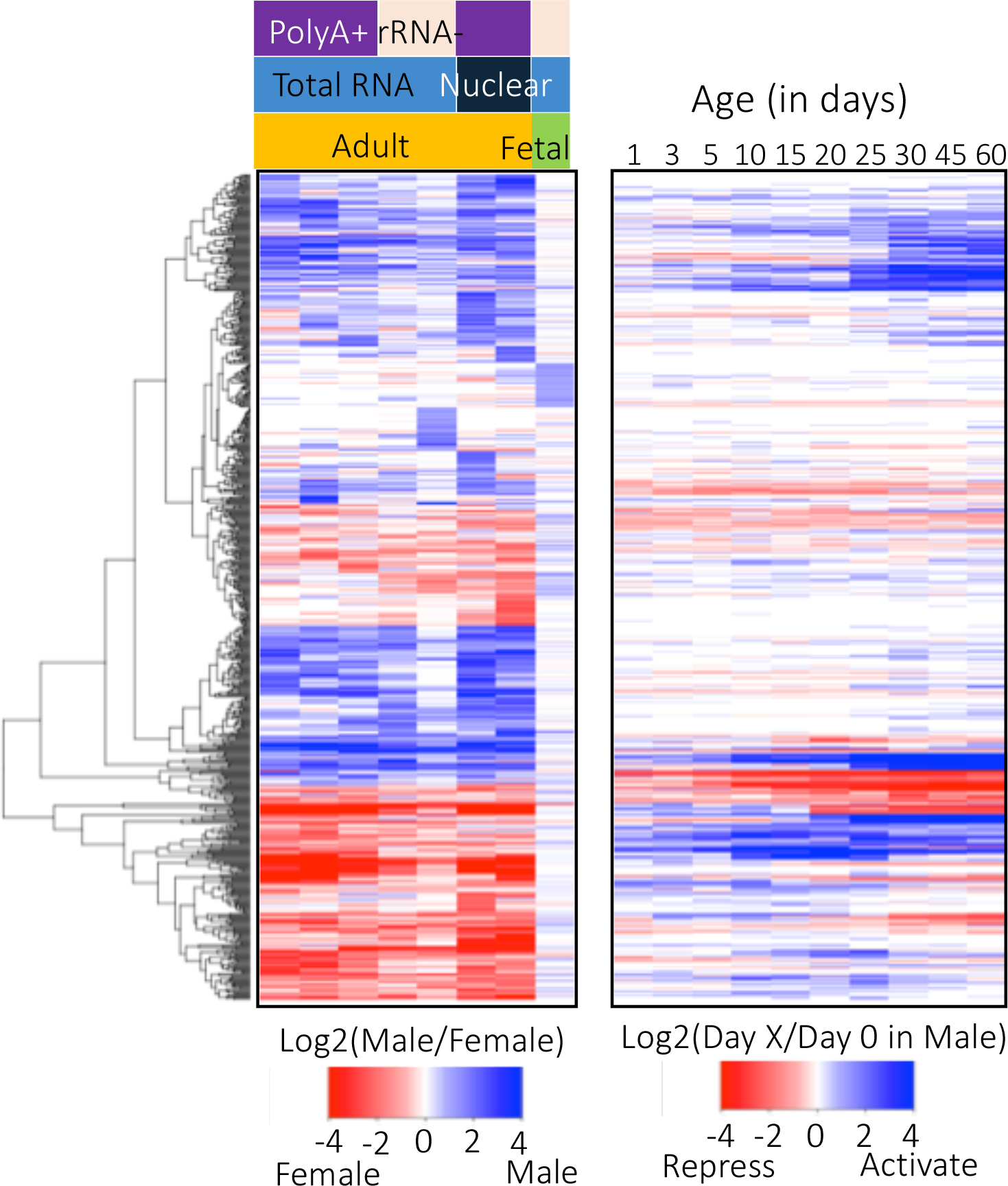
A Heatmap of 684 sex-specific lncRNAs in different RNA-Seq datasets. The first 8 columns in the heatmap correspond to the 8 datasets used to identify sex-specific lncRNAs; blue indicates male-specific gene expression, and red indicates female-specific gene expression. The last 10 columns track the changes of gene expression in male liver at the ages specified at top, as compared to birth (day 0). Blue denotes up regulation of gene expression, compared to day 0, and red denotes down regulation of gene expression at the specified age, as compared to day 0.

**Figure S4.**
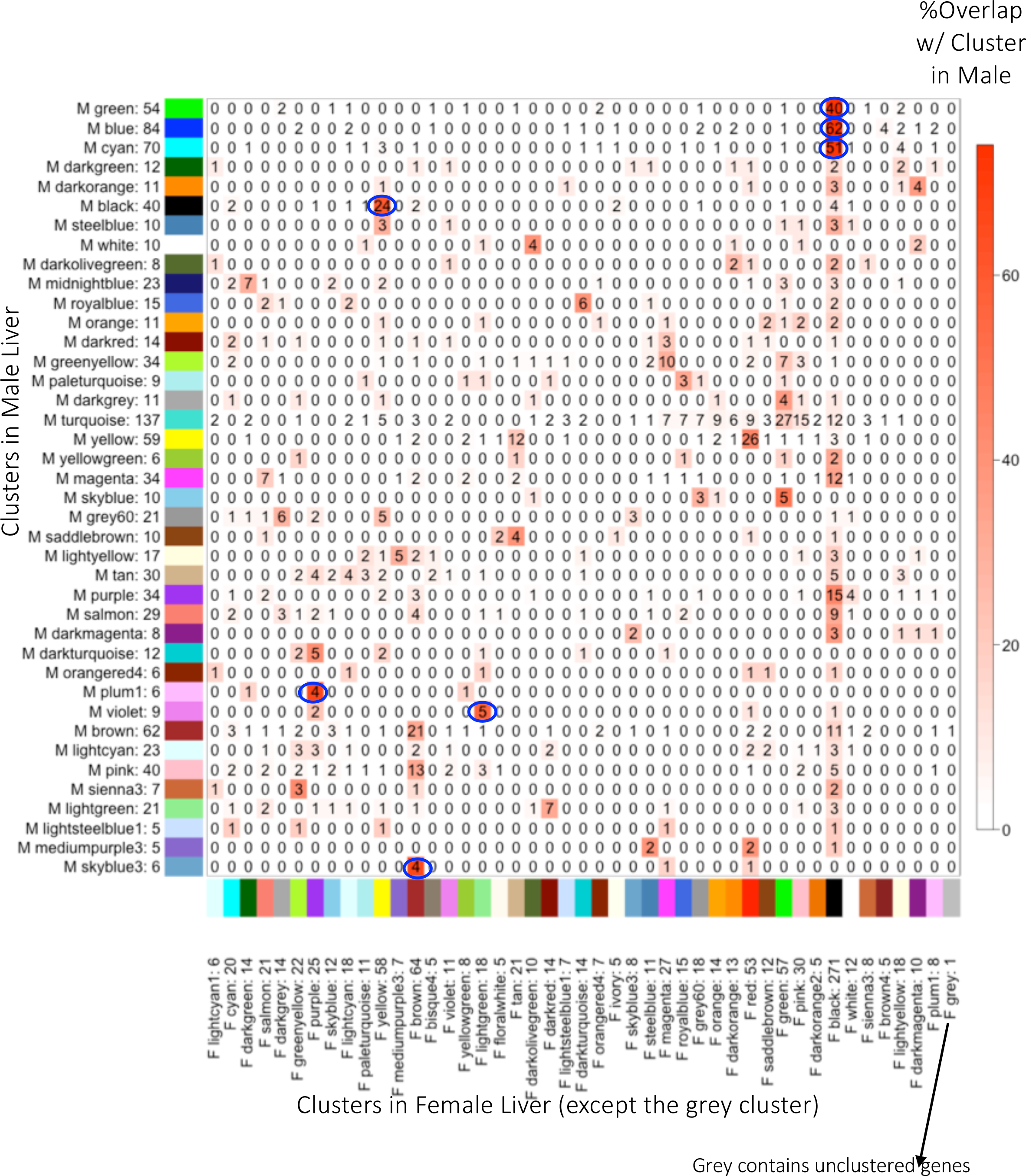
Overlap between clusters discovered in male *vs*. female liver. The number of genes in each matrix cell represents the number of genes in the cluster discovered in male DO mouse liver (y-axis) that are present in the indicated clusters discovered in female DO mouse liver (x-axis). The last cluster in female: grey, is not considered a cluster; it contains 1 gene that was not able to be clustered in female liver. The color represents the percentage of overlap with respect to the cluster discovered in male liver. The number written along each axis after each cluster name is the number of genes in each cluster. Circles denote strong overlap, where >50% of gene members in a specific cluster in male (row) are in a specific cluster in female (column). Also, see Fig 5.

**Figure S5.**
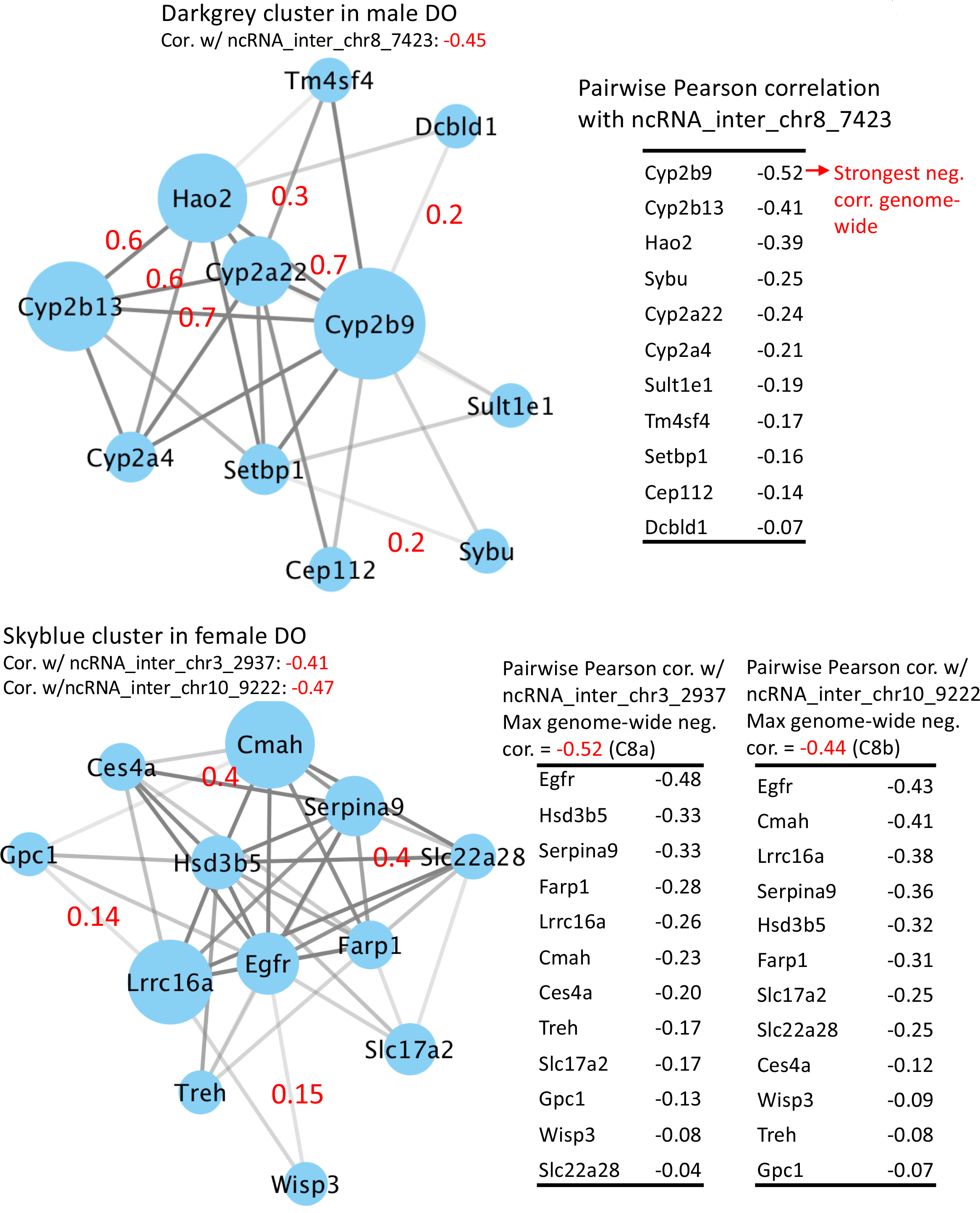
Examples of gene modules that are negatively correlated with sex-specific lncRNAs (Table 2). Nodes denote genes, and edges denote the correlation between genes. The darkness of the edges indicates the correlation strength between genes. The size of each node represents the cumulative gene correlations for a gene, where the larger the node is, the more high correlations involve that node. Graphs were drawn using Cytoscape [76]. Pairwise Pearson correlation for select edges are shown in red. Pairwise correlations between gene members with the lncRNA with which the respective module is correlated are tabulated for select gene modules.

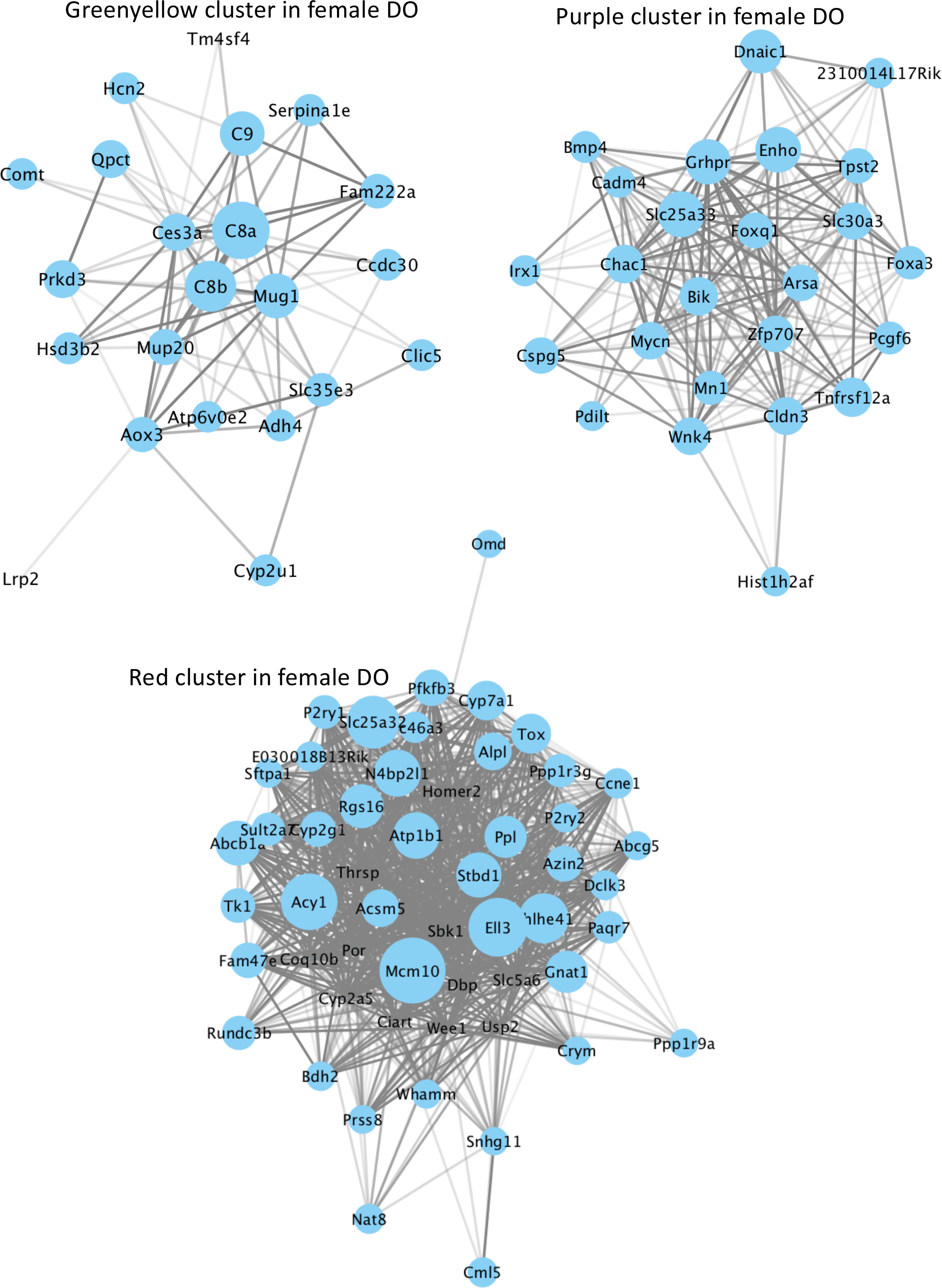

## Notes

This work was supported in part by NIH grant DK033765 (to DJW)

